# Mild SARS-CoV-2 infection results in long-lasting microbiota instability

**DOI:** 10.1101/2022.12.07.519508

**Authors:** Vaibhav Upadhyay, Rahul Suryawanshi, Preston Tasoff, Maria McCavitt-Malvido, G. Renuka Kumar, Victoria Wong Murray, Cecilia Noecker, Jordan E. Bisanz, Yulin Hswen, Connie Ha, Bharat Sreekumar, Irene P. Chen, Susan V Lynch, Melanie Ott, Sulggi Lee, Peter J. Turnbaugh

**Affiliations:** Department of Microbiology and Immunology, G.W. Hooper Research Foundation, University of California, San Francisco, CA 94143, USA; Department of Medicine, University of California San Francisco, University of California, San Francisco, CA 94143, USA; Benioff Center for Microbiome Medicine, Department of Medicine, University of California, San Francisco, CA 94143, USA; Gladstone Institutes, San Francisco, CA 94158 USA; Department of Epidemiology and Biostatistics and the Bakar Computational Health Institute at the University of California San Francisco, San Francisco, CA 94158 USA; Department of Pediatrics, University of California San Francisco, University of California, San Francisco, CA 94143, USA; Chan Zuckerberg Biohub, San Francisco, CA 94158

**Keywords:** COVID-19, SARS-CoV-2, non-hospitalized patients, human gut microbiome, gastrointestinal symptoms, microbial ecology

## Abstract

Viruses targeting mammalian cells can indirectly alter the gut microbiota, potentially compounding their phenotypic effects. Multiple studies have observed a disrupted gut microbiota in severe cases of SARS-CoV-2 infection that require hospitalization. Yet, despite demographic shifts in disease severity resulting in a large and continuing burden of non-hospitalized infections, we still know very little about the impact of mild SARS-CoV-2 infection on the gut microbiota in the outpatient setting. To address this knowledge gap, we longitudinally sampled 14 SARS-CoV-2 positive subjects who remained outpatient and 4 household controls. SARS-CoV-2 cases exhibited a significantly less stable gut microbiota relative to controls, as long as 154 days after their positive test. These results were confirmed and extended in the K18-hACE2 mouse model, which is susceptible to SARS-CoV-2 infection. All of the tested SARS-CoV-2 variants significantly disrupted the mouse gut microbiota, including USA-WA1/2020 (the original variant detected in the United States), Delta, and Omicron. Surprisingly, despite the fact that the Omicron variant caused the least severe symptoms in mice, it destabilized the gut microbiota and led to a significant depletion in *Akkermansia muciniphila*. Furthermore, exposure of wild-type C57BL/6J mice to SARS-CoV-2 disrupted the gut microbiota in the absence of severe lung pathology.

**IMPORTANCE:** Taken together, our results demonstrate that even mild cases of SARS-CoV-2 can disrupt gut microbial ecology. Our findings in non-hospitalized individuals are consistent with studies of hospitalized patients, in that reproducible shifts in gut microbial taxonomic abundance in response to SARS-CoV-2 have been difficult to identify. Instead, we report a long-lasting instability in the gut microbiota. Surprisingly, our mouse experiments revealed an impact of the Omicron variant, despite producing the least severe symptoms in genetically susceptible mice, suggesting that despite the continued evolution of SARS-CoV-2 it has retained its ability to perturb the intestinal mucosa. These results will hopefully renew efforts to study the mechanisms through which Omicron and future SARS-CoV-2 variants alter gastrointestinal physiology, while also considering the potentially broad consequences of SARS-CoV-2-induced microbiota instability for host health and disease.

## INTRODUCTION

Mammalian viruses exhibit bidirectional interactions with the gut microbiota (the trillions of microorganisms colonizing the gastrointestinal tract) (1). The gut microbiota and its aggregate gene content (the gut microbiome) contributes to protective immunity from influenza (2, 3) and respiratory syncytial virus (4), whereas human immunodeficiency virus (HIV) is associated with marked perturbations in gut microbial community structure and function (5). The gut microbiota is also distinctive in patients with severe SARS-CoV-2 infections requiring hospitalization relative to healthy controls (6–8); however, the direct causal effects of SARS-CoV-2 relative to concomitant changes in host immunity, diet, and pharmacotherapy remain unknown. Decreased bacterial richness is a reproducible marker of SARS-CoV-2 infection (7–10), whereas the specific bacterial taxa correlated with infection varies between studies, in part due to differences in the built environment (10). Furthermore, the generalizability of these findings to subjects with milder cases that do not require hospitalization and have less severe symptoms is unclear.

The impact of SARS-CoV-2 in the outpatient setting is a timely question as current estimates suggest the vast majority (92%) of individuals in the United States that test positive for SARS-CoV-2 will not require hospitalization (11). The predominant variant at the time of this manuscript is Omicron (12), which is less likely to require hospitalization (13), aided in part by pre-existing immunity due to vaccination and prior waves of infection (14). Despite these encouraging trends, a growing number of non-hospitalized adults still develop long-lasting symptoms persisting months after clearing the virus (15, 16), highlighting the importance of understanding the mechanisms responsible. Considered in light of emerging evidence that the microbiome can exhibit “ecological memory” of past events (17), we hypothesized that even mild cases of SARS-CoV-2 could still disrupt the gut microbiota, potentially contributing to phenotypes months later.

Here, we present an analysis of subjects participating in the COVID-19 Host Immune Response Pathogenesis (CHIRP) study. CHIRP was an exploratory study of primarily outpatients and their household contacts collected between May and August 2020. Prior work on this cohort revealed SARS-CoV-2 specific CD8^+^ T cells are maintained well into convalescence (recovery from disease), even in mild disease (18, 19). However, these prior studies did not analyze the gut microbiota. To address this knowledge gap, we used paired 16S rRNA gene and metagenomic sequencing from longitudinal samples collected from CHIRP cases and household controls. We confirmed and extended these findings using two mouse models of SARS-CoV-2: the K18-humanized Angiotensin Converting Enzyme 2 (K18-hACE2) mouse model (20) and C57BL/6J mice. SARS-CoV-2 binds to the human ACE2 receptor but some variants cannot interact with the orthologous mouse protein (21, 22). The K18-hACE2 mouse expresses human ACE2 under the Keratin-18 promoter, leading to expression in lung epithelium, and provides an experimentally tractable model to study multiple SARS-CoV-2 variants (23). In contrast, C57BL/6J mice develop infection without meaningful lung pathology with a subset of variants (24) and present a complementary mouse model of infection. Taken together, our results define a significant and long-lasting impact of mild SARS-CoV-2 infection on the gut microbiota.

## RESULTS

### Lack of a reproducible shift in the gut microbiota following mild cases of SARS-CoV-2

We evaluated the gut microbiota of outpatients with SARS-CoV-2 infections during the first year of the pandemic. Samples were collected in the weeks to months after initial infection (maximum 154 days after initial positive test results; **Fig. S1**). 53 longitudinal stool samples were collected from 18 subjects enrolled in CHIRP, including 6 men and 12 women whose ages ranged from 19-71 years in a Case-Control household study design (**Table S1**). DNA was extracted from samples and used for paired 16S rRNA gene sequencing (16S-Seq) and metagenomic sequencing (MGS). We generated 135,600±7,236 (mean±sem) high-quality 16S rRNA gene reads/sample and 54.0±2.19 million (mean±sem) high-quality MGS reads/sample (**Table S2A**).

On average, the gut microbiomes of SARS-CoV-2 Cases were similar to Controls. Both groups were primarily colonized by members of the *Firmicutes, Bacteroidota*, and *Actinobacteriota* phyla across the sampling period (16S-Seq data; **Fig. 1A**). PERMANOVA testing adjusted for longitudinal sampling did not reveal a significant difference in gut microbial community structure (*p*_adj_=1) (**Fig. 1B**). Similarly, there were no changes in bacterial diversity (**Fig. S2A**) or 16S-Seq ASV abundances, which were highly correlated between groups (**Fig. 1C**). The structure of MGS data at the pathway-(**Fig. S2B**) and gene-level (**Fig. S2C**) was similar in Cases and Controls. Pathway-(**Fig. 1D**) and gene families (**Fig. 1E**) were highly correlated between Cases and Controls. Statistical testing confirmed the lack of a reproducible shift in phylum, ASV, or metabolic pathway levels (*p*_adj_>0.1, see *Methods*).

**FIG. 1.**
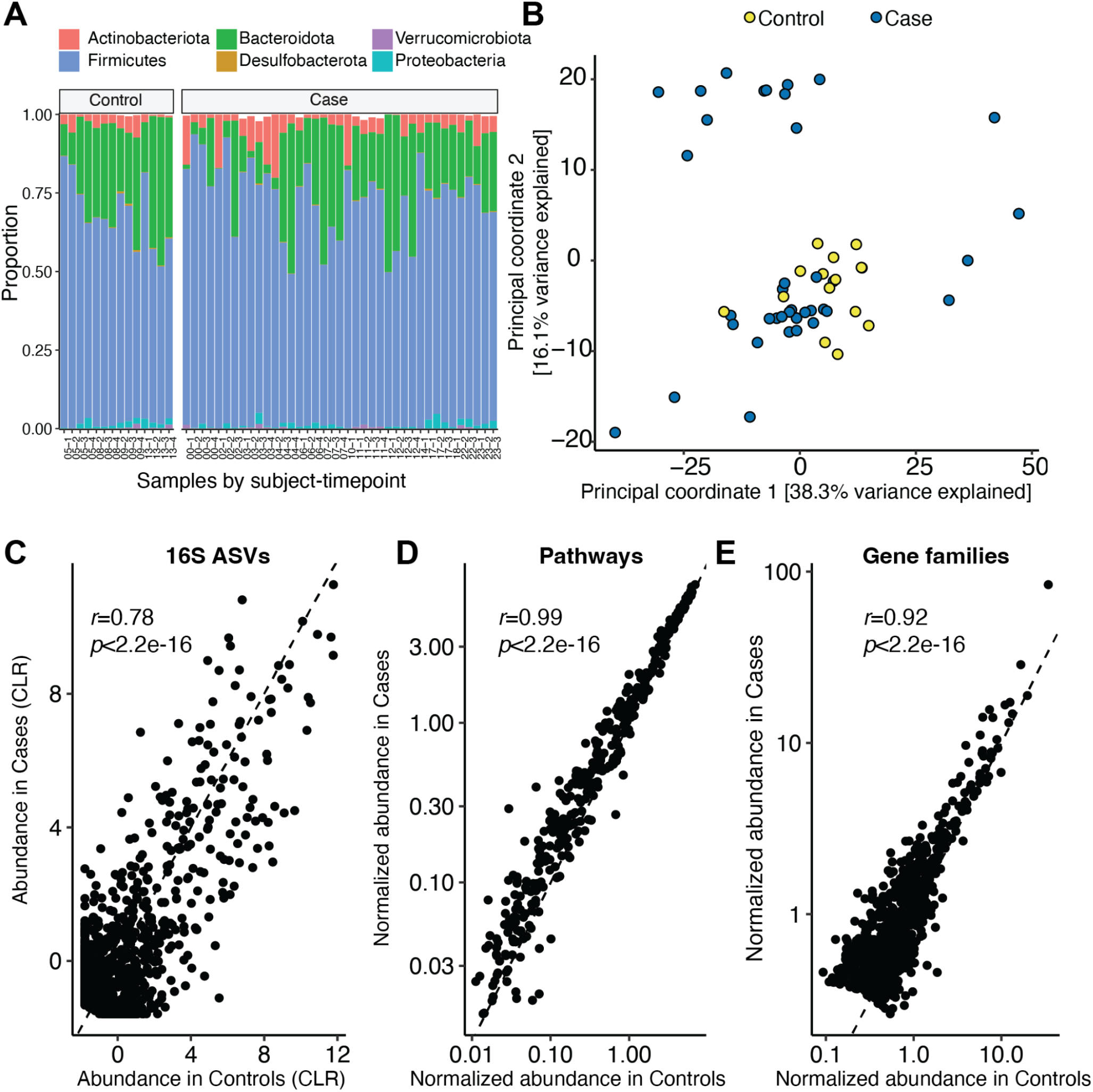
Mild SARS-CoV-2 infection does not reproducibly change the gut microbiome months after infection. **(A)** Phylum-level taxonomic summary of 16S-Seq data for Cases and Controls. **(B)** Principal coordinate analysis of 16S-Seq data **(C-E)** Scatter plots reveal a significant correlation between Cases and Controls for the abundance of **(C)** 16S-Seq ASVs, **(D)** MGS pathways, and **(E)** MGS gene families. Data represents 53 samples spanning 18 subjects (14 Cases, 4 Controls).

### Long-lasting microbiota instability following SARS-CoV-2 infection

We hypothesized that the lack of specific differences in the gut microbiomes of Cases and Controls may have been due to high levels of variation among the SARS-CoV-2 infected individuals. Consistent with this hypothesis, visualization of our 16S-Seq and MGS data by Principal Coordinates Analysis (PCoA) and quantification of β-dispersion (25) both demonstrated a marked and significant increase in temporal variation of the gut microbiotas of Cases relative to Controls (**Figs. 2A-B**). Samples from Cases deviated across sampling timepoints more than Controls in terms of 16S-Seq ASV abundance (**Fig. 2C**) and MGS functional pathways (**Fig. 2D**). We calculated the coefficient of variation (*CV*), a statistical measure of dispersion, for all ASVs or MGS functional pathway feature ranks. The *CV* of features for Cases was significantly higher for taxonomic information inferred by 16S-Seq (**Fig. 2E**) and functional information inferred by MGS sequencing (**Fig. 2F**) relative to Controls. *CV* was negatively correlated with feature abundance for both datasets in Cases and Controls (**Fig. S3**).

**FIG. 2.**
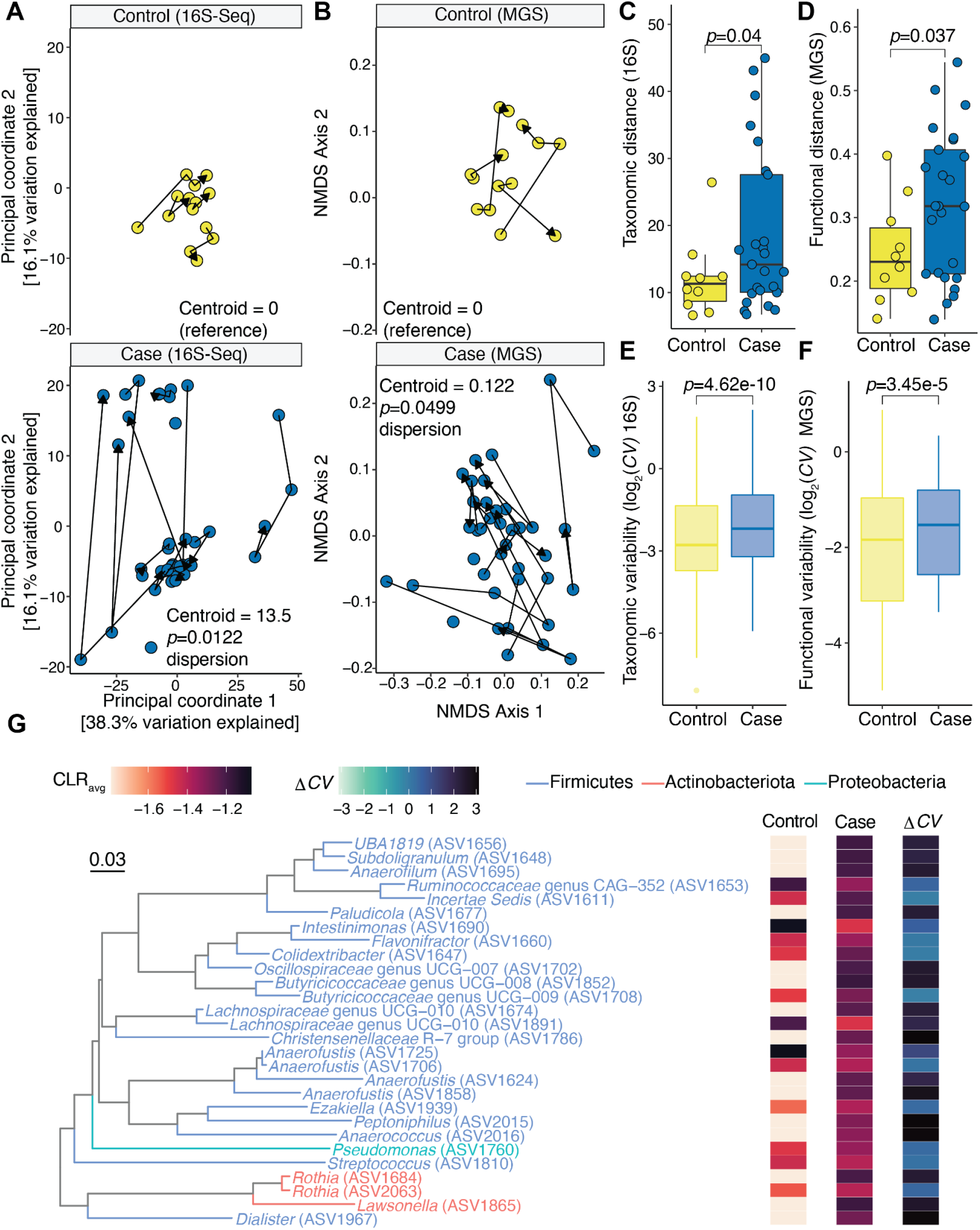
The human gut microbiota is less stable following mild SARS-CoV-2 infection. **(A-B)** Principal coordinate analysis of 16S-Seq data **(A)** or NMDS of MGS data **(B)**. A given subject is connected by a line with arrows indicating successive sample collection; *p*-value for β-dispersion from the vegan package for SARS-CoV-2 positivity is displayed and is calculated between Cases and Controls and adjusted using Tukey’s multiple comparisons. **(C-D)** Subsequent distances between successive points using different data sets as shown in for 16S-Seq from **Fig. 2A (C)** or MGS from **Fig. 2B (D)** for a given subject are plotted and grouped by Cases or Controls. Student’s *t*-test between groups is annotated. **(E-F)** *CV* was calculated for each feature making up 16S-Seq **(E)** or MGS **(F)**. A Dunn’s post test of Kruskal-Wallis ANOVA is annotated. **(G)** The ASVs with the highest *CV* (>97.5% percentile) are plotted with CLR abundance and *ΔCV* displayed where *ΔCV* = *CV*_Case_-*CV*_Control_. n=18 subjects, 53 samples.

Next, we sought to identify which bacterial species were most variable following SARS-CoV-2 infection, and visualized the most variable ASVs (**Fig. 2G**). Low abundance organisms were among the most variable species as determined by *CV* (**Fig. 2G**). Variable bacteria spanned distantly related phyla (**Fig. 2G)**, and included *Rothia* species which have been associated with SARS-CoV-2 and correlated with differences in the built environment (9).

Complementary analyses of our metagenomic data confirmed these trends (**Fig. S4**). Gene family abundance exhibited significantly more variability (β-dispersion metric) in Cases relative to Controls (**Fig. S4A**). Gene families trended towards more variation between sample points for Cases compared to Controls (**Fig. S4B**). Gene family variability assessed by *CV* was also significantly higher in Cases relative to Controls (**Fig. S4C**).

Taken together, these data indicate that the human gut microbiome can be destabilized months after initial infection with SARS-CoV-2. However, it is not possible to infer a causal role of SARS-CoV-2 infection in destabilizing the gut microbiota based only on observational studies in humans given the clear potential for confounding factors. By focusing on outpatient sampling, we were able to rule out the confounding effects of hospitalization and treatment in prior studies (26); however, our longitudinal samples corresponded to a period of extensive social distancing (**Fig. S5**) that could have feasibly impacted the gut microbiota (27). Thus, we turned to an established mouse model of SARS-CoV-2 infection, wherein environmental and genetic variables could be controlled to test the causal role of viral infection in shaping the mouse gut microbiota.

### SARS-CoV-2 alters the gut microbiota in susceptible mice

We tested the impact of 3 SARS-CoV-2 variants on gut microbiota in K18-hACE2 mice: WA1, Delta, and Omicron (BA.1). These variants differ in their phenotypic impacts in K18-hACE2 mice, with each successive temporal SARS-CoV-2 variant resulting in milder infection than the prior variant (23). 50 K18-hACE2 female mice were housed in an Animal Biosafety Level 3 (ABSL3) facility and longitudinal stool samples were collected following infection and successive planned sacrifice for virological assessment of gut and lung tissues (**Fig. S6A and Table S2B**). As described (23), lung viral titers were significantly different between variants (**Fig. S6B**), with WA1 having the highest titer and Omicron the lowest titer. The SARS-CoV-2 mRNA transcripts for envelope (E) and nucleocapsid (N) genes were detectable using nucleic acid amplification within small intestinal tissue (**Figs. S6C-D**) with trends mirroring those observed for viral plaque assays from the lung (**Fig. S6B**). To evaluate the gut microbiota, we performed 16S-Seq on 55 samples, resulting in 435,539±7,099 (mean±sem) high-quality reads/sample (**Table S2B**).

All SARS-CoV-2 variants led to a dramatic and significant impact on the mouse gut microbiota. Visualization of all three variants compared to the Uninfected mice showed clearly distinct patterns of grouping (**Fig. 3A**); however, they did not reach statistical significance by PERMANOVA testing after adjusting for longitudinal sampling. Bacterial diversity increased over time following infection with all three variants (**Fig. 3B**). We observed differences at the phylum level for four phyla between groups (two-way ANOVA for *Verrucomicrobiota, p*=6.09e-8; *Firmicutes, p*=1.29e-5; *Bacteroidota, p*=9.08e-4; and *Proteobacteria, p*=1.10e-2). There was a striking loss of *Verrucomicrobiota* in Omicron infected mice and a subtle reduction in *Proteobacteria* in WA1 infected mice, with reciprocal shifts between *Firmicutes* and *Bacteroidota* in these groups (**Fig. 3C**). Notably, *Akkermansia* diminished over time in response to infection with WA1, Delta, and Omicron (**Fig. 3D**).

**FIG. 3.**
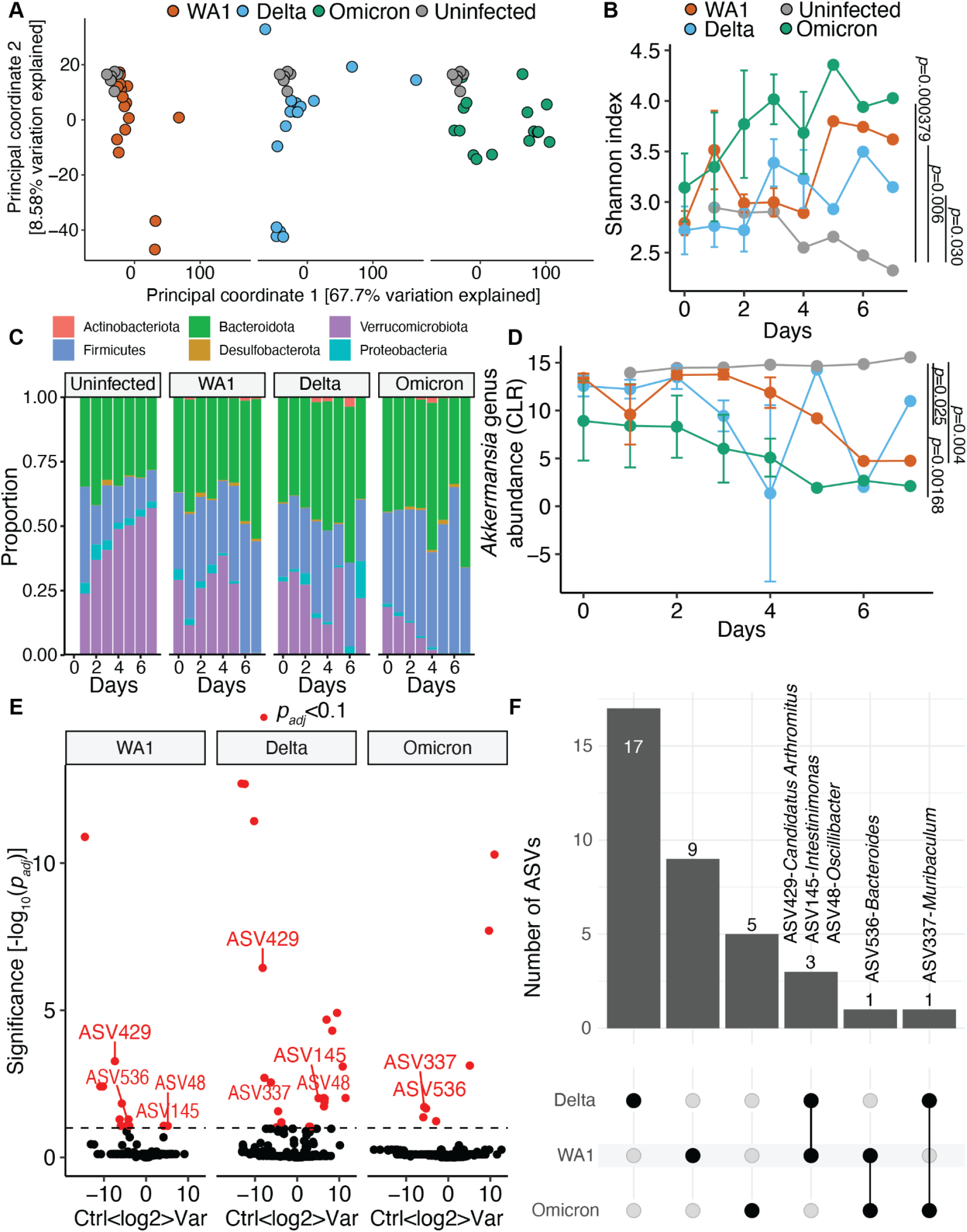
SARS-CoV-2 variants have significant and heterogeneous impact on the mouse gut microbiota. **(A)** Principal coordinate analysis of 16S-Seq data for mouse samples with the Uninfected group and each of the SARS-CoV-2 variants. **(B)** Shannon index for 16S-Seq data colored by variant of concern against the days after inoculation on the x-axis (Days). **(C)** Phylum-level taxonomic summaries for each group plotted against days after inoculation of infected groups (Days). **(D)** *Akkermansia* CLR for each variant by day of sampling is plotted. **(B, D)** *p*-values displayed are results of two way ANOVA and reflect values comparing each variant to the Uninfected group. **(E-F)** A linear mixed effect model was created for each variant and **(E)** a Volcano plot of adjusted *p*-values versus difference in CLR abundance is displayed and **(F)** features that met an Benjamini Hochberg adjusted *p*-value<0.1 with overlap between groups is displayed in columns. Features impacted by more than one variant are annotated in both **(E)** and **(F)**. n=10 unique cages with 55 total samples from 50 mice.

Given the multi-strain design of our experiment, we sought to understand whether the impact of SARS-CoV-2 on the gut microbiota was variant-specific. We analyzed the ASV data using a linear mixed effect model to identify shared and unique responses to each strain (**Fig. 3E**). For each variant, most ASV changes were unique to the variant itself (**Fig. 3F** and **Table S3A**). There were 5 ASVs that were impacted by two strains with no ASVs consistently impacted by all three strains (**Fig. 3E-F**). Taken together, these results indicate that the impact of SARS-CoV-2 varies between strains and does not track with overall disease severity.

Finally, we tested if SARS-CoV-2 infection of the K18-hACE2 model would recapitulate the microbiota destabilization phenotype that we observed in humans. We generated rarefaction curves from the longitudinally sampled mice in a given cage. Rarefaction curves provided a clear indication of microbiota instability in infected mice, with stereotyped curves in Uninfected controls and a massive spread in all 3 SARS-CoV-2 infected groups (**Fig. 4A**). The range of ASVs detected was 84-95 (controls), 70-320 (WA1), 53-249 (Delta), and 84-324 (Omicron). Microbiota instability was also clear by PCoA (**Fig. 4B**), with a significant increase in β-dispersion for Delta and Omicron. Both changes in the gut microbial variation (**Fig. 4C**) and species variability (**Fig. 4D**) were significantly increased following infection. Similar to humans, there was a negative correlation between *CV* and ASV abundance (**Fig. 4E**). In Uninfected mice, the correlation between *CV* and abundance was nearly bimodal with features having a Centered-Log-Ratio (CLR) value<0 having high *CV* values compared to those with mean CLR values above 0. In contrast, during infection there was significantly increased *CV* across organisms of all abundances, though the negative correlation between *CV* and ASV abundance was still preserved (**Fig. 4E**). Inspection of the most variable ASVs revealed low abundance features (**Fig. 4F**) that produced no reads in Uninfected mice. Similar to our data from human subjects, these features spanned distantly-related species. Taken together, these findings support a causal and strain-specific role of SARS-CoV-2 in microbiota instability.

**FIG. 4.**
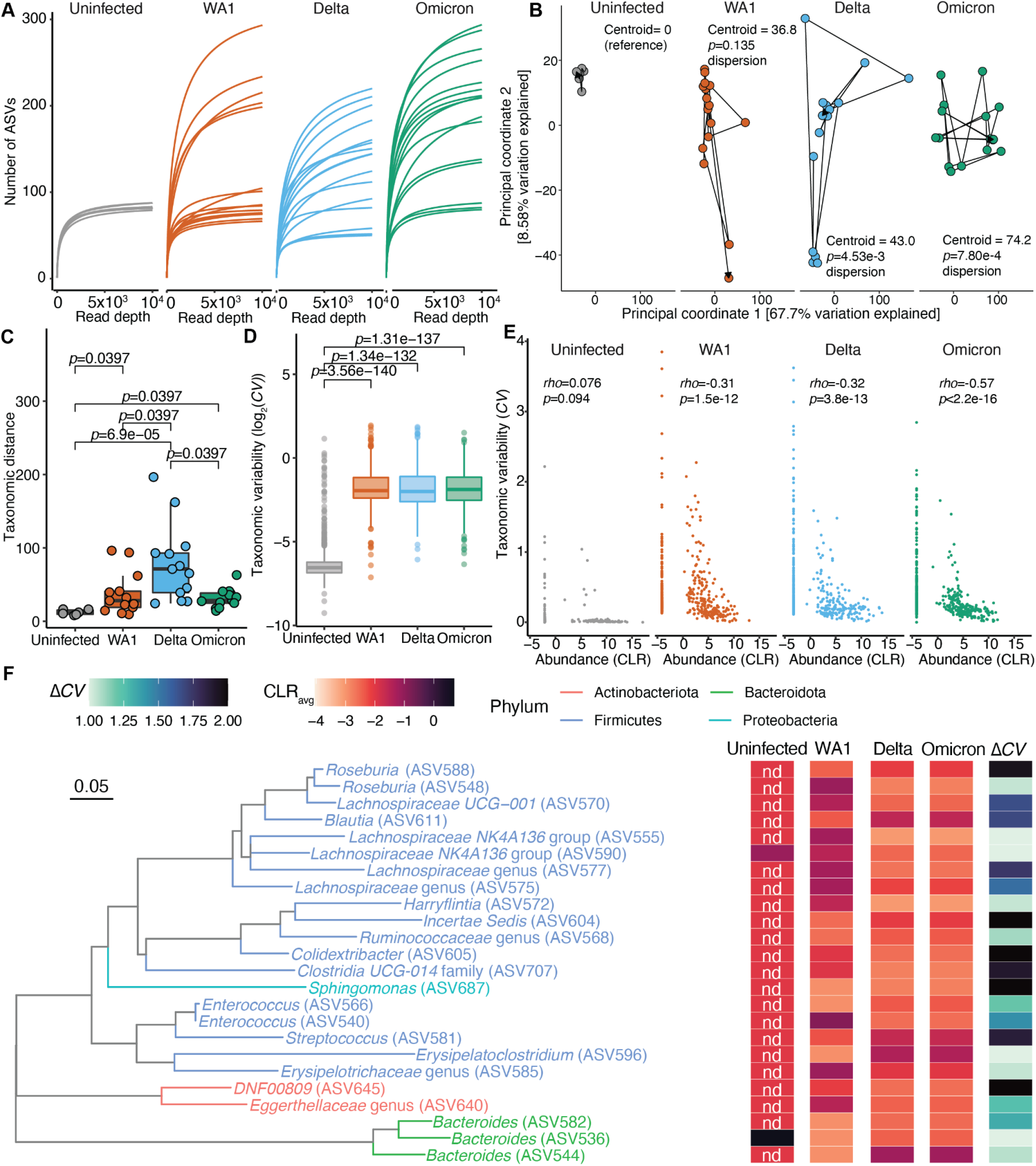
SARS-CoV-2 destabilizes the mouse gut microbiota. **(A)** Rarefaction curves for Uninfected, WA1, Delta, or Omicron infected mice are plotted for all samples from each group. **(B)** Principal coordinates 1 and 2 of 16S-Seq data for all analyzed samples derived from the same distance matrix and separated by group to facilitate visualization. Points are connected sequentially by day after infection from a given cage. The distance from the Uninfected reference group’s centroid is displayed for the right three panels. The *p*-value for β-dispersion from the vegan package for COVID Status is displayed and compares the variant to the Uninfected reference group and adjusted using Tukey’s multiple comparisons. **(C)** Subsequent distances between successive points in **(B)** for a given cage are plotted and grouped by SARS-CoV-2 variants of concern. **(D)** *CV* for each 16S feature is plotted by SARS-CoV-2 variants of concern. **(C-D)** Dunn’s post-hoc test of Kruskal-Wallis analysis of variance is displayed. **(E)** A correlation between Taxonomic variability and Abundance is displayed and a Spearman’s correlation coefficient superimposed for each group. **(F)** ASVs with the highest *CV* (>97.5% percentile) are plotted with CLR abundance. For the Uninfected group, ASVs that had 0 reads present are annotated as not detected (nd). n=10 unique cages with 55 total samples from 50 mice.

### SARS-CoV-2 impacts the gut microbiota in the absence of lung pathology

Given the robust shift in the gut microbiota in response to Omicron, the mildest variant in the K18-hACE2 model, we hypothesized that the immunological or other types of host responses to viral inoculation could alter the gut microbiota independent of host disease. To test this hypothesis, we turned to wild-type C57BL/6J mice that are resistant to severe lung pathology from SARS-CoV-2 infection (24). We infected C57BL/6J mice with Beta and Omicron variants and longitudinally collected stool samples for 20 days (**Fig. S6E**). We performed 16S-Seq on 33 samples, resulting in 398,412±10,679 (mean±sem) high-quality reads/sample (**Table S2C**). Live virus was detectable in the lungs at day 2 post infection; however, we did not detect any live virus in the gastrointestinal tract (**Fig. S6F**).

Remarkably, Beta and Omicron variants resulted in a pronounced shift in the gut microbiota of C57BL/6J infected mice by day 2 post infection (**Fig. 5A**). Similar to K18-hACE2 mice, the adjusted PERMANOVA was not significant. Beta infected mice showed greater bacterial diversity compared to Uninfected mice, although Omicron infected mice C57BL/6J mice did not show this trend (**Fig. S7A**). Similar to K18-hACE2 infected mice, phylum-level abundance was significantly altered in response to infection (two-way ANOVA for *Verrucomicrobiota, p*=6.97e-8; *Firmicutes, p*=1.54e-8; *Bacteroidota, p*=1.10e-2; and *Proteobacteria, p*=1.07e-6). *Proteobacteria* were depleted in Beta and Omicron infected mice, though not as marked or pronounced as the depletion of *Verrucomicrobiota* in these same groups (**Fig. 5B**). Correspondingly, *Akkermansia* were depleted in both Beta and Omicron infected groups (**Fig. 5C**). Numerous ASVs were identified as being depleted or elevated by both Beta and Omicron, with 11 ASVs overlapping between both groups (**Fig. 5D-E** and **Table S3B**). Finally, while there was no difference in β-dispersion or distance traveled between sampling points for Beta or Omicron infected mice compared to the Uninfected group (**Fig. S7B-C**), the variability of taxonomic features was significantly greater in mice infected with Beta or Omicron (**Fig. 5F**).

**FIG. 5.**
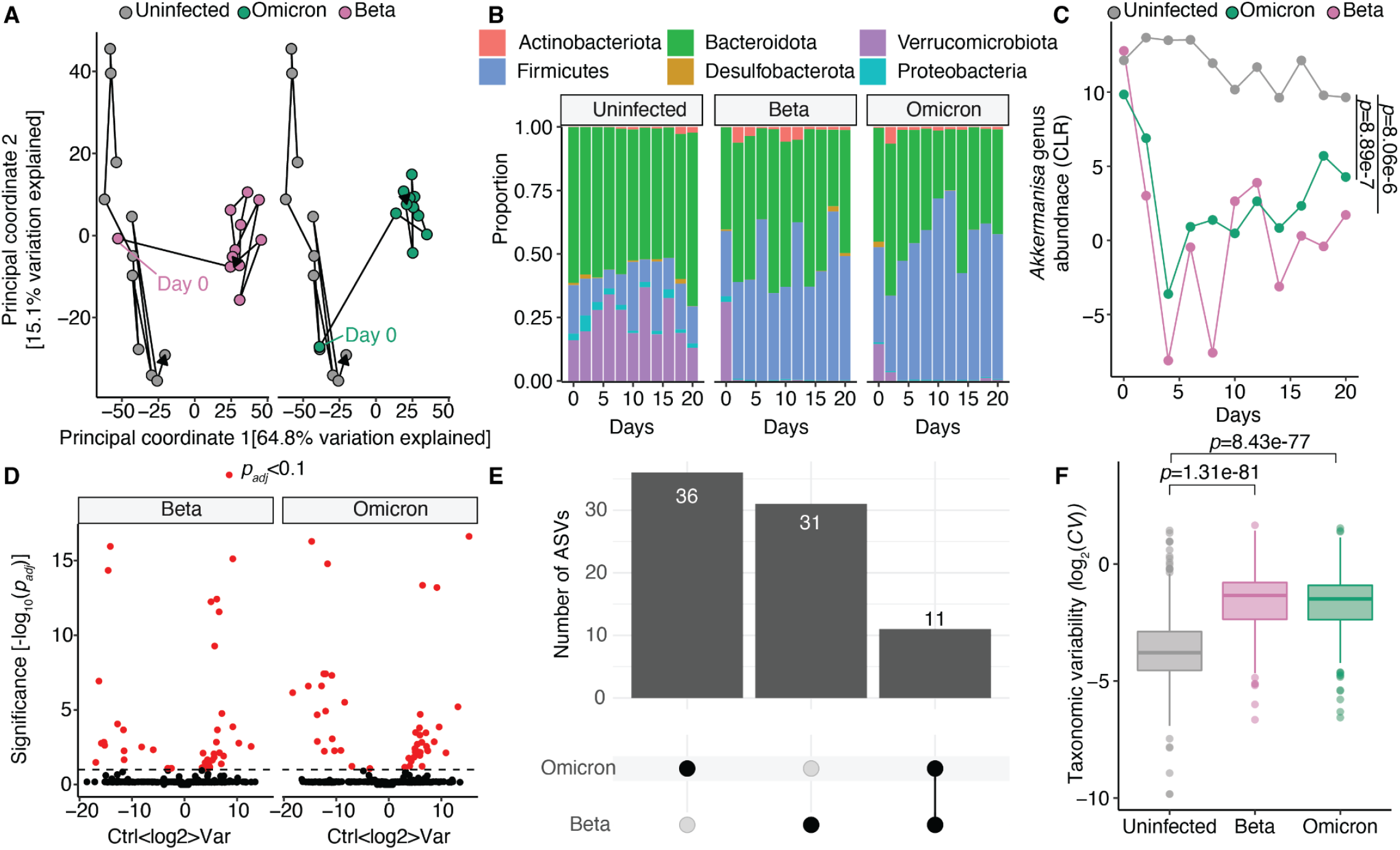
Mild infection in wild-type mice result in unique destabilization to the gut microbiota and loss of *Akkermansia*. **(A)** Principal coordinate analysis of C57BL/6J mice comparing communities after Beta and Omicron variant infection are shown relative to Uninfected mice. **(B)** Phylum level taxonomic summaries for Uninfected, Beta or Omicron infected mice sampled from days 0 to 20 after inoculation (Days) are displayed. In **(C)** *Akkermansia* genus CLR for each variant is plotted against day of sampling post-inoculation on the x-axis (Days). *p*-values displayed are results of two way ANOVA and reflect values comparing each variant to the Uninfected group. **(D-E)** A linear mixed effects model was created for each variant and **(D)** a Volcano plot of adjusted *p*-values versus difference in CLR abundance is displayed and in **(E)** the number of ASVs shared between variants is shown. **(F)** Taxonomic variability is plotted comparing Uninfected, Beta, and Omicron infected mice. n=15 mice with 5 mice per group in 3 cages and 33 total longitudinally collected samples.

## DISCUSSION

Our results in humans and mice demonstrate that the gut microbiota is destabilized following mild cases of SARS-CoV-2 infection; however, the cellular and molecular mechanisms responsible remain to be elucidated. SARS-CoV-2 most likely impacts the gut microbiome through effects on host immune or epithelial cells (28). However, despite its minimal impact on the host, Omicron still led to a dramatic collapse of the mucin-dependent gut Verrucomicrobium *A. muciniphila*. These results may suggest that SARS-CoV-2 can lead to dysfunction of the intestinal goblet cells, due to either direct infection or the increased level of intestinal cytokines. While early work on WA1 suggested that it cannot bind to goblet cells (29), this work conflicts with recent data demonstrating goblet cell hyperplasia in response to severe SARS-CoV-2 infection (28). The impact of Omicron on *A. muciniphila* was notable and suggests the continued evolution of SARS-CoV-2 may have preserved its ability to perturb the intestinal mucosa.

The downstream consequences of microbiota instability for COVID-19 pathophysiology will be important to assess. Persistent symptoms have been observed in a subset of adults characterized principally by fatigue, headache, anosmia, and dyspnea though impacting every organ system (16). A recent report showed findings similar to ours in that taxonomic changes in the gut microbiome were captured as late as 6 months after SARS-CoV-2 infection and were linked to prolonged symptoms, though instability of the gut microbiome was not assessed (30). Gastrointestinal symptoms such as nausea, abdominal pain or diarrhea, may be exacerbated by more recently evolved variants either through their direct impact on the intestine or resulting destabilization of the gut microbiome caused by infection.

It is also important to consider the impact of disruptions in the gut microbiota to host responses in other contexts. SARS-CoV-2 significantly alters the pulmonary immune response (23). *A. muciniphila* has established links to promoting activation of exhausted T cells (31), and eliciting high quantity mucosal IgA responses (32). Disrupting *A. muciniphila* or other bacterial species that control immune function could be important as the host balances viral clearance with tissue damage that might lead to outcomes like acute lung injury or prolonged symptoms after infection (33).

Importantly, we discovered a marked instability in the human gut microbiota following SARS-CoV-2 infection, complicating efforts to identify bacterial genes, pathways, and taxonomic groups that consistently differentiated Cases and Controls in humans. Across all data types and groups in our study, instability was the hallmark of SARS-CoV-2 infection. This data is in line with a concept that has been referred to as the “Anna Karenina Principle”, wherein disease-associated microbial communities are distinct from the microbiotas of healthy individuals but lack shared features (34). Similar observations have been made in the context of broad-spectrum antibiotics, autoimmunity, and enteric bacterial infections (34); however, this is to our knowledge the first evidence that mild cases of a respiratory virus infection can lead to microbial community-wide instability in the gastrointestinal tract. Longitudinal studies of other common viral infections in the outpatient setting will be important in order to test the generalizability of these findings, coupled to larger cohorts of SARS-CoV-2 patients. Furthermore, the individualized nature of the microbiota’s response to infection highlights the potential for incorporating microbiome signatures into precision medicine.

Our study has several limitations. Although the CHIRP samples analyzed in this study are a vanishingly rare commodity of SARS-CoV-2 naïve individuals following initial infection, the number of subjects was small and each individual was unevenly sampled. In addition, we had limited metadata regarding infection-associated changes in diet or other lifestyle factors that could compound microbiome susceptibility, including the social distancing practices of each participant. The duration and scope of our mouse studies were limited due to the logistical challenges of performing experiments under ABSL3 conditions.

Despite these limitations, our data clearly shows that the gut microbiota can be destabilized, at least in some individuals, following mild SARS-CoV-2 infection. These changes were independent of disease severity and distinct between SARS-CoV-2 variants, potentially due to variant-specific effects on goblet cell function and mucosal integrity. It is notable that SARS-CoV-2 has reached virtually every corner of the world (35). While trends towards milder infections are a cause for celebration, it will be important to consider that mild infections have the potential for months-long destabilization of the gut microbiota and that more recently evolved variants still have a pronounced impact on the gut microbiota in mouse models. These results will be important to revisit and evaluate in the future as we approach the next era of the pandemic and move toward understanding the long-term impact of prior SARS-CoV-2 infection for our health and the health of our microbial co-conspirators.

## MATERIALS AND METHODS

### CHIRP subject enrollment

Subjects were recruited to the CHIRP study and provided informed consent (IRB 20-30588). Subjects were asked to produce a nasal PCR indicating SARS-CoV-2 status at the time of enrollment. Subjects underwent survey based questionnaires at visits and provided stool samples which were frozen at -80°C. One participant (CHIRP-4108) had symptoms but did not have a positive PCR test at the time of study enrollment, with subsequent research based PCR testing as inconclusive. CHIRP-4108 is the child of CHIRP-4107 and CHIRP-4109. Two subjects, CHIRP-4100 and CHIRP 4106 (a Case and Control from separate households) reported antibiotic exposure prior to study enrollment. For the purposes of this study, only subjects with both a definitive positive PCR swab and symptoms are referred to as Cases and all other individuals are Controls. Household relationships are listed in **Table S1**.

### Mouse SARS-CoV-2 infection models

Protocols for animal use were approved (AN169239-01C) by the Institutional Animal Care and Use committees at the University of California, San Francisco and Gladstone Institutes. Mice were housed in a temperature-controlled specific pathogen-free facility with a 12-h light–dark cycle and *ad libitum* access to water and laboratory rodent chow. The study involved intranasal infection of 10^4^ plaque-forming units (PFU) of 6–8-week old female K18-hACE2 mice with Delta, Omicron, and WA1 variants of SARS-CoV-2. In the case of C57BL/6J mice, mice were infected with 10^3^ PFU of Beta or Omicron variants of SARS-CoV-2. All stool samples from a cage of mice were pooled into one tube with DNA/RNA shield 1-5 cages of infected mice sampled per time point. Where possible, two of the aggregated samples were selected for sequencing otherwise the resulting slurry was sequenced. In the case of K18-hACE2 mice, a full cage was euthanized at 2, 4 and 7 days post-infection for lung viral titer assessment or if meeting criteria for sacrifice on days 5 and 6 as described in **Fig. S5**. The lung and gut tissues were analyzed for viral titer using plaque assays as described in (23).

### Stool DNA extraction

Stool DNA was extracted using a modified protocol within a BSL2 biological safety cabinet given subjects had known infection with SARS-CoV-2. Samples were handled on dry ice and containers were decontaminated before and after use with 70% ethanol. Briefly, samples were extracted using phenol-chloroform and 5% hexadecyltrimethylammonium bromide. Samples underwent two rounds of bead-beating for 45 seconds at a rate of 5.5 m/s and underwent heat denaturation for 15 minutes at 65°C. Polyethylene glycol was used to precipitate DNA, which was then washed in 70% ethanol. This method has been described previously (36). For mouse stool samples, roughly two stool pellets were selected from the DNA/RNA shield mixture and processed for 16S-Seq as described below.

### 16S rRNA gene amplicon sequencing

DNA was amplified using Kapa-HiFi Hotstart (KK2502, Kapa Biosystems) using primers to 16S-V4 regions (V4-515F - TCGTCGGCAGCGTCAGATGTGTATAAGAGACAGGTGCCAGCMGCCGCGGTAA, V4-806R - GTCTCGTGGGCTCGGAGATGTGTATAAGAGACAGGGACTACHVGGGTWTCTAAT) on a BioRad CFX 384 real-time PCR instrument with four serial 10 fold dilutions of extracted DNA template. Individual sample dilutions in the exponential phase were selected using an OpenTrons OT2 for subsequent indexing PCR using a dual GoLay error correcting index primers (37). DNA concentration was measured using a PicoGreen assay (P7589, Life Technologies) and samples were pooled at equimolar concentrations. Agencourt AMPure XP magnetic beads were used to purify the pooled PCR product, and the samples were subsequently sequencing on an Illumina MiSeq using 15% PhiX spiked in for sequencing. Mouse samples were amplified using V4 primers as previously described (38). Amplicon reactions were pooled at equimolar concentrations and purified using the Agencourt AMPure XP magnetic beads. The pooled library was loaded onto the Illumina NextSeq 550 platform using 40% PhiX spiked in for sequencing.

### 16S rRNA gene sequencing analysis

Primer sequences were trimmed to 220 base pairs for forward reads and 150 base pairs for reverse reads and, where necessary, adapters were trimmed using the cutadapt plugin in Qiime2. DNA sequences underwent quality filtering, denoising, chimera filtering utilizing DADA2 as implemented in QIIME2 (39). Taxonomy was assigned to Amplicon Sequence Variants (ASVs) using the SILVA v138 database (40). Two negative control samples were processed in this manner. Greater than 98% of both negative controls reflected chimeric reads and were ∼ 10^2^ lower in final read content than the lowest positive control. Negative control samples were subsequently removed from analysis. ASVs were filtered for those found with at least 10 reads in 3 samples and subsampled to even sampling depth using MicrobeR (41).

Samples underwent a PhILR transformation using the *de novo* generated phylogenetic made using the philr package (42). β-dispersion was assessed in this distance matrix using the function beta.disper in the R package vegan with resulting *p*-values were adjusted using Tukey’s multiple comparison test, and PERMANOVA testing was conducted using the adonis2 function with with the following formula (DM ∼ Household+COVID_Status) (25). For PERMANOVA testing, the adonis2 command from the vegan package (25) was used and blocks of permutation were set to 10,000 and restricted by individual subject using the how function in base R using participant ID. For **Fig. 2**, distances for individuals from which successive measurements were available were selected from the larger PhILR distance matrix and initial measurements (e.g. 0 distance) were excluded. A linear mixed effect model with the following formula (Center Log Ratio (CLR) ∼ COVID_Status+ (1|Subject ID) was iterated over all ASVs using lmerTest for **Fig. 1** and a model fitting CLR ∼ Variant + Day_Post_Infection + (1|Cage) was fit for **Fig. 3, Fig. 5** and **Table S3A-B** (43). ASVs for whom the model failed due to singularity were excluded from the analysis. Resulting *p*-values were corrected using the p.adjust command in R and the Benjamini-Hochberg correction. Calculation of coefficients of variance (*CV*) were completed by transforming CLR data to ranks and then dividing the mean of a feature by its standard deviation using the group by command and on a per sample basis iterated across all features for 16S-Seq, taxonomic MGS, and functional MGS data. In **Fig. 4F**, a CLR value was assigned to all indicated ASVs in the control groups and upon reviewing the raw read counts, there were 0 reads in all Uninfected samples for all ASVs identified in **Fig. 4F** as indicated in the figure and legend.

For mouse 16S data, Day 0 time points were excluded from PERMANOVA testing and linear mixed effects models for variant infected mice as these timepoints reflected variant infected mice prior to infection. PERMANOVA testing was restricted by longitudinally sampled cages by using the how function in base R and 10,000 permutations. Differences in *Akkermansia* were determined by comparing all ASVs assigned to the genus *Akkermansia* between groups after CLR transformation of genus ASVs. Two-way anova was calculated using the anova_test tool in the rstatix package, where the indicated dependent variable in the figure was a function of two independent variables (e.g. SARS-CoV-2 variant and days post infection).

### Metagenomic sequencing

Extracted DNA was prepared using a Nextera XT DNA library preparation kit from Illumina and methods were followed as specified by the manufacturer. 18 out of 58 samples lacked sufficient concentration to create 50 ng of sample material, and for these 20-30 uL of samples were used to provide the maximum amount of template material at tagmentation and PCR. Samples were amplified using 6 cycles of PCR using the IDT Illumina Type B indexing primers. Sample DNA concentrations were estimated using a PicoGreen assay (P7589, Life Technologies), pooled to equimolar concentrations using an OpenTrons OT2, and cleaned using the Agentcourt AMPure XP magnetic beads. The resulting library was sequenced on an Illumina NovaSeq instrument at UCSF’s Center for Advanced Technology.

### Metagenomic analyses

Shotgun libraries were processed using Humann3 with unstratified pathway data or gene families as outputs (44). Normalized abundance was calculated as Abundance normalized by Genome Equivalents as estimated by Microbecenssus (45). Histograms of feature counts by number of samples present were created and a cut off of feature presence in 3 samples was used for pathway data or gene-family data. While all gene-family data trimmed in this manner was used to construct distance matrices and related analyses, low abundant features were trimmed and only the top 20,191 gene families were evaluated for associations with SARS-CoV-2 using a linear mixed effect model and represented in **Fig. 1E**. Pathway and gene family data was normalized to reads per kilobase million (RPKM)*10^3^ cutoffs of greater than 0.0005. A Canberra distance matrix was calculated for filtered functional data with subsequent PERMANOVA testing utilizing adonis2, and nMDS plots were created using the metaMDS command (25). PERMANOVA testing, dispersion assessment, and distances between points were done as described for 16S-Seq.

### SARS-CoV-2 E and N protein quantitative PCR

Mice were infected with the indicated variants of SARS-CoV-2 and RNA was extracted from small intestinal tissue that underwent hommogenization. Quantitative Polymerase Chain Reaction (qPCR) was conducted using SYBR Green. qPCR was conducted for SARS-CoV-2 E and N Genes (E Gene Forward primer: 5′-ACAGGTACGTTAATAGTTAATAGCGT-3′; Reverse primer: 5′-ATATTGCAGCAGTACGCACACA-3’, N Gene Forward primer: 5′-AAATTTTGGGGACCAGGAAC-3′; Reverse primer: 5′-TGGCACCTGTGTAGGTCAAC-3’) and normalized to Gapdh gene (Forward primer: 5’-AGGTCGGTGTGAACGGATTTG*-3*’); Reverse primer: 5’-TGTAGACCATGTAGTTGAGGTCA-3’). Reactions were 10 μL and conducted in 384-well plates using an annealing temperature of 60°C on a CFX384 Touch Real-Time PCR Detection System (Bio-Rad).

### Social distancing variable measurements

We obtained population level social distancing variables from Cubiq. Cubiq provided two pieces of data for each zip code and date combination. The Cubiq Mobility Index (CMI) quantifies movement in users in a given region per day. Movement was calculated using a derivative factor indicating the distance between opposite corners within a box around locations for users on a given day. CMI values can be interpreted as follows (5-100,000 meters; 4-10,000 meters; 3-1000 meters; 2-100 meters, 1-10 meters). The “Home Percentage” variable is the estimated number of users sheltering-in-place, which is calculated by the number of users moving less than 330 feet from home on a given day.

### Statistical analysis

Data was analyzed either in this R version 4.0.4, or an enterprise version of R studio 3.5.1 for 16S-Seq data and R version 3.6.1 for shotgun data on the Wynton Computing cluster (a co-op based computing cluster at UCSF). Unless otherwise stated, data was analyzed using the following software packages in R: tidyverse, ggplot2, and rstatix (46, 47).

### Data Availability

All sequencing data pertaining to this manuscript has been uploaded to the NIH Sequence Read Archive under the accession number PRJNA885137 (MGS) and PRJNA907010 (16S-Seq). MGS data was depleted of human reads prior to upload. Metadata for each sample is included in the SRA deposited samples and corresponds to identified sample names within the supplementary tables of this manuscript.

## SUPPLEMENTAL MATERIAL

Supplemental material is available online only.

Fig S1, PDF file, 231 KB.

Fig S2, PDF file, 495 KB.

Fig S3, PDF file, 803 KB.

Fig S4, PDF file, 5.3 MB.

Fig S5, PDF file, 174 KB.

Fig S6, PDF file, 1.2MB.

Fig S7, PDF file, 482 KB.

Tables S1-S3, XLSX file, 124 KB.

## ACKNOWLEDGEMENTS

Funding was provided by the National Institutes of Health (R01HL122593, R01AT011117, R01DK114034, P.J.T., and F31AI164671 I.P.C.) and the Benioff Center for Microbiome Medicine (BCMM 2013260 V.U.). P.J.T. is a Chan Zuckerberg Biohub Investigator and held an Investigators in the Pathogenesis of Infectious Disease Award from the Burroughs Wellcome Fund. M.O received support from the James B. Pendleton Charitable Trust, Roddenberry Foundation, P. and E. Taft, and Gladstone Institutes.

Conceptualization, V.U., S.L., P.J.T.; Data curation, V.U.; Formal Analysis, V.U.; Funding acquisition, M.O., S.L., P.J.T.; Investigation, V.U., R.S., P.T., M.M., R.K., V.M., B.S., I.P.C., Y.H.; Methodology, C.N.; Project administration, P.J.T.; Resources, P.J.T.; Software, C.N., J.E.B.; Supervision, C.H., S.V.L., M.O., S.L., P.J.T.; Validation, V.U., P.J.T.; Visualization, V.U.; Writing – original draft, V.U.; Writing – review & editing, P.J.T.

P.J.T. is on the scientific advisory boards for Pendulum, Seed, and SNIPRbiome; there is no direct overlap between the current study and these consulting duties. The other authors declare no competing interests.

## SUPPLEMENTAL FIGURES AND LEGENDS

**FIG. S1.**
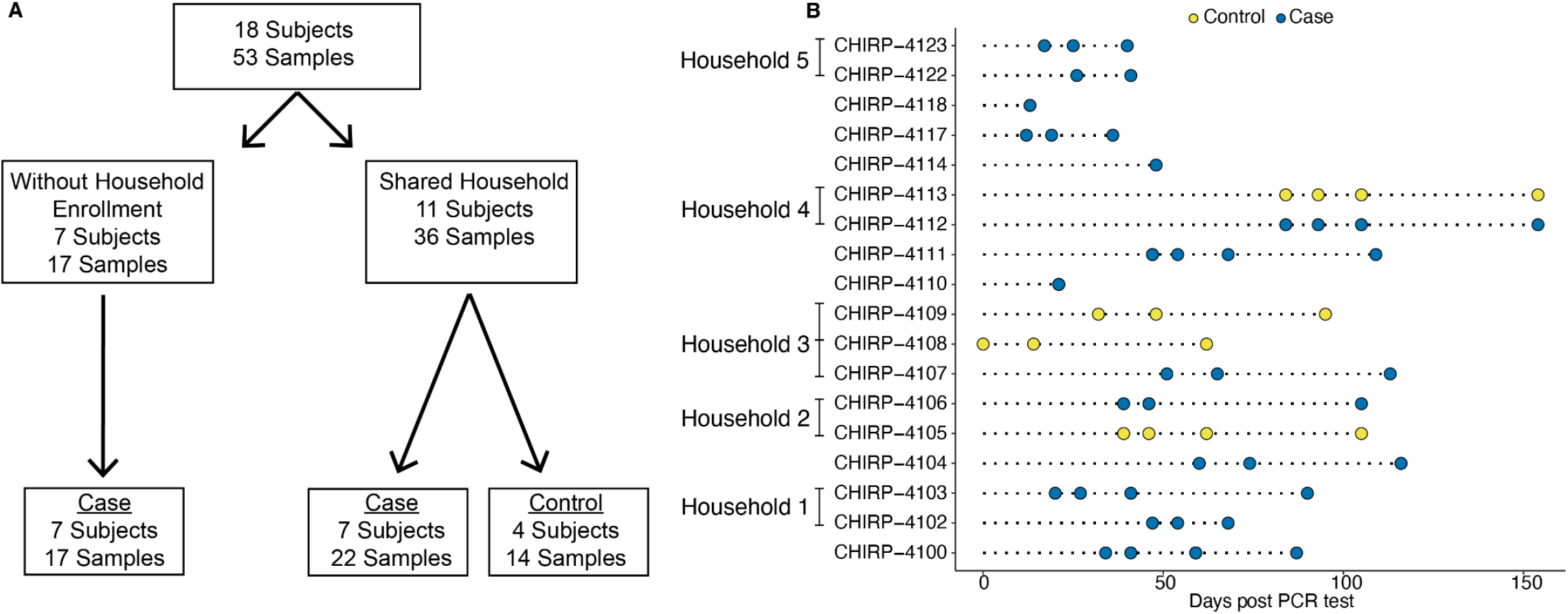
Clinical study sampling scheme. **(A)** CONSORT diagram for the overall CHIRP study and the microbiome sub-analysis. **(B)** Sample collection with respect to positive test result for SARS-CoV-2 subject in household. Brackets indicate individuals from the same household as described in **Supplementary Table 1**.

**FIG. S2.**
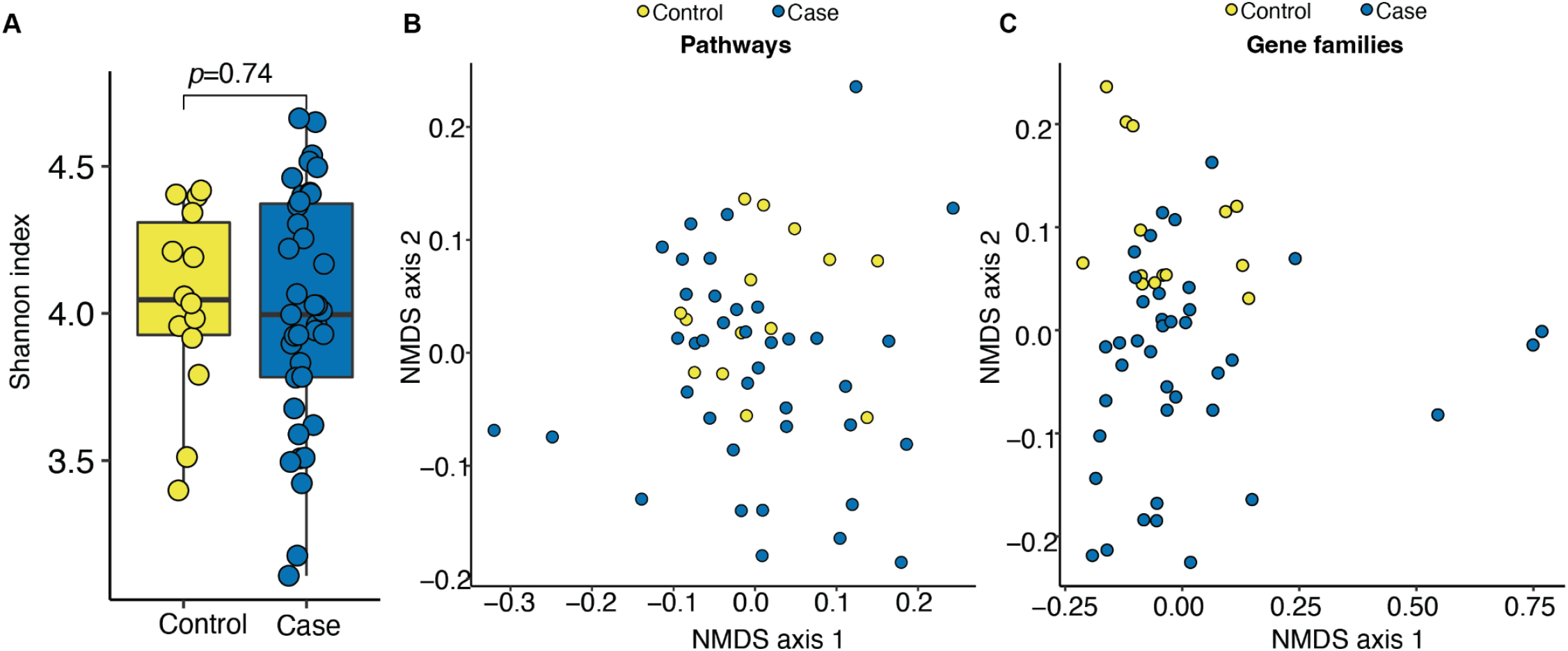
Lack of a consistent signature of mild SARS-CoV-2 infection based on 16S-seq and functional metagenomic data. Panel **(A)** reflects 16S-Seq data and **(B-C)** are MGS data. **(A)** Microbial diversity was comparable between groups based on the Shannon Index. *p*-value, Student’s t-test. **(B-C)** Humann3 unstratified pathway data **(B)** or gene families **(C)** from MGS show that overall community structure was similar between groups, visualized by PCoA, analogous to 16S-Seq data (PERMANOVA *p*_adj_=1 for both pathway and gene family data). n=18 subjects, 53 samples.

**FIG. S3.**
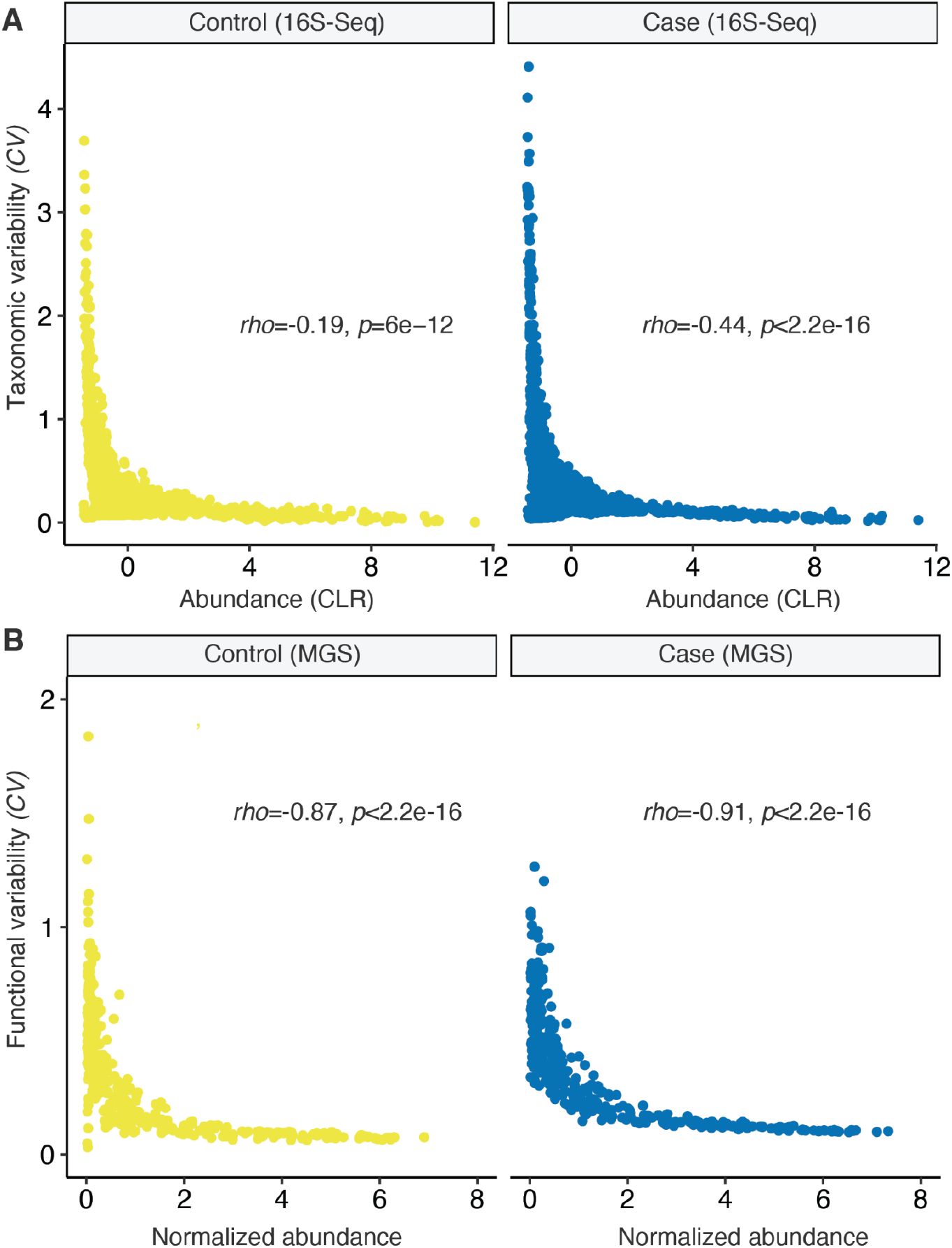
Feature variability negatively correlates with abundance in Cases and Controls. The *CV* of 16S-Seq ASVs **(A)** or MGS Pathways **(B)** was plotted against the abundance of each and separated between Cases and Controls. A Spearman’s correlation coefficient is annotated indicating a negative correlation between *CV* and feature abundance in all groups. n=18 subjects, 53 samples.

**FIG. S4.**
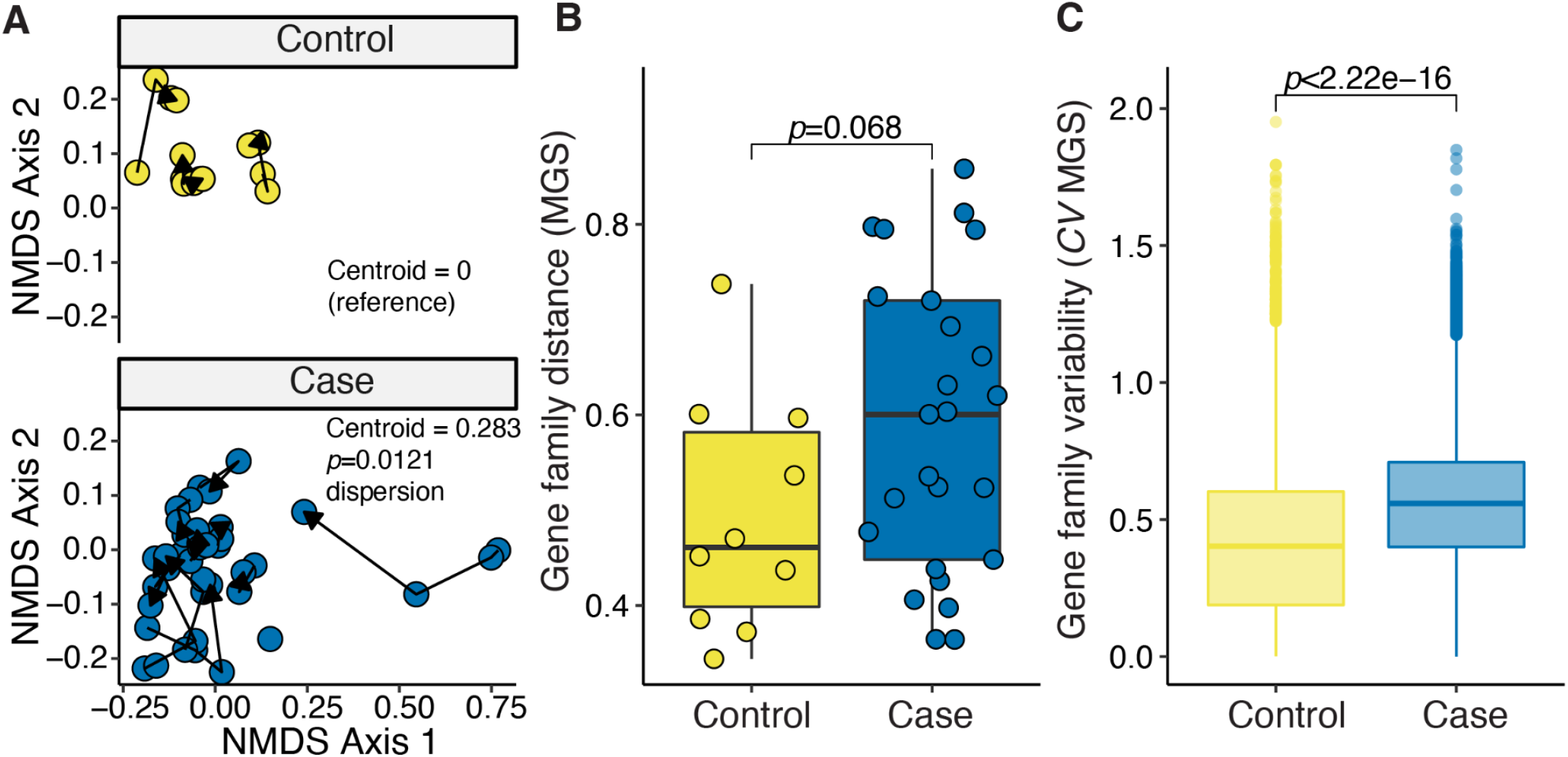
The human gut microbiome is less stable following mild SARS-CoV-2 infection as measured by metagenomic data annotated to gene families. **(A)** Principal coordinate analysis of MGS data annotated to gene families. A given subject is connected by a line with arrows indicating successive sample collection; *p*-value for β-dispersion from the vegan package for SARS-CoV-2 positivity is displayed and is calculated between Cases and Controls and adjusted using Tukey’s multiple comparisons. **(B)** Subsequent distances between successive points as shown in **Fig. S4A**. Student’s *t*-test between groups is annotated. **(C)** *CV* was calculated for each feature making up gene family MGS data. A Dunn’s post test of Kruskal-Wallis ANOVA is annotated. n=18 subjects, 53 samples.

**FIG. S5.**
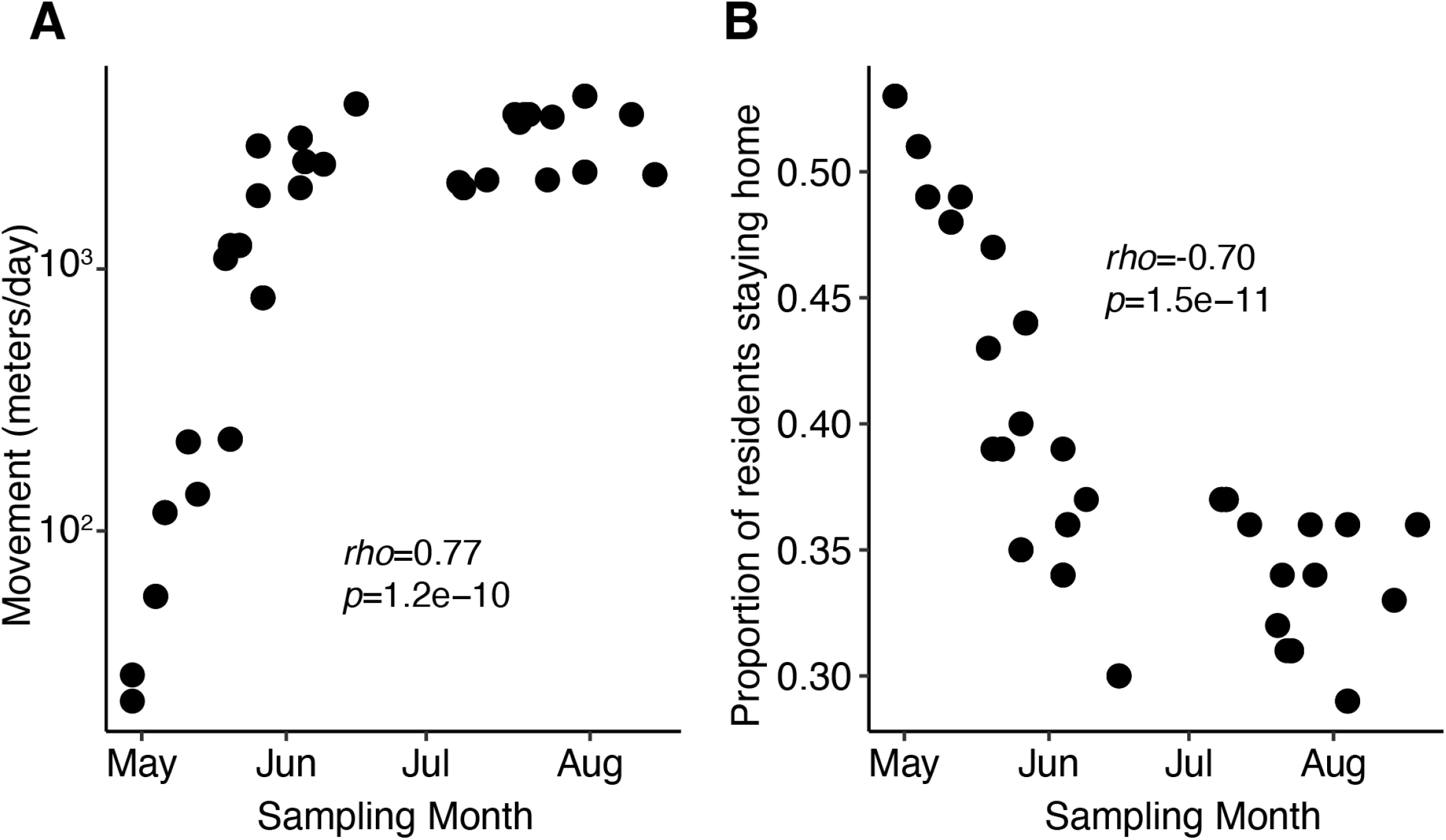
Variation in population level social distancing practices for samples at the time of collection. Population level social distancing data was obtained from the Cubiq organization which utilizes cellular telephone movement of users to approximate movement and number of residents remaining at home. This was done for each sample based on the zipcode of the CHIRP participant and the date of sample collection. **(A)** The Cubiq Mobility Index is plotted on the y-axis and approximately reflects movement in 10^n^ meters/day where n is Cubiq Mobility Index **(B)** The proportion of residents estimated to reside at home per day by Cubiq is plotted on the y-axis. In both **(A)** and **(B)** the values from Cubiq are plotted against the month in 2020 of sampling on the x-axis. A Spearman’s correlation coefficient and *p*-value is plotted. n=16 subjects, 48 samples for which this type of data was available from Cubiq.

**FIG. S6.**
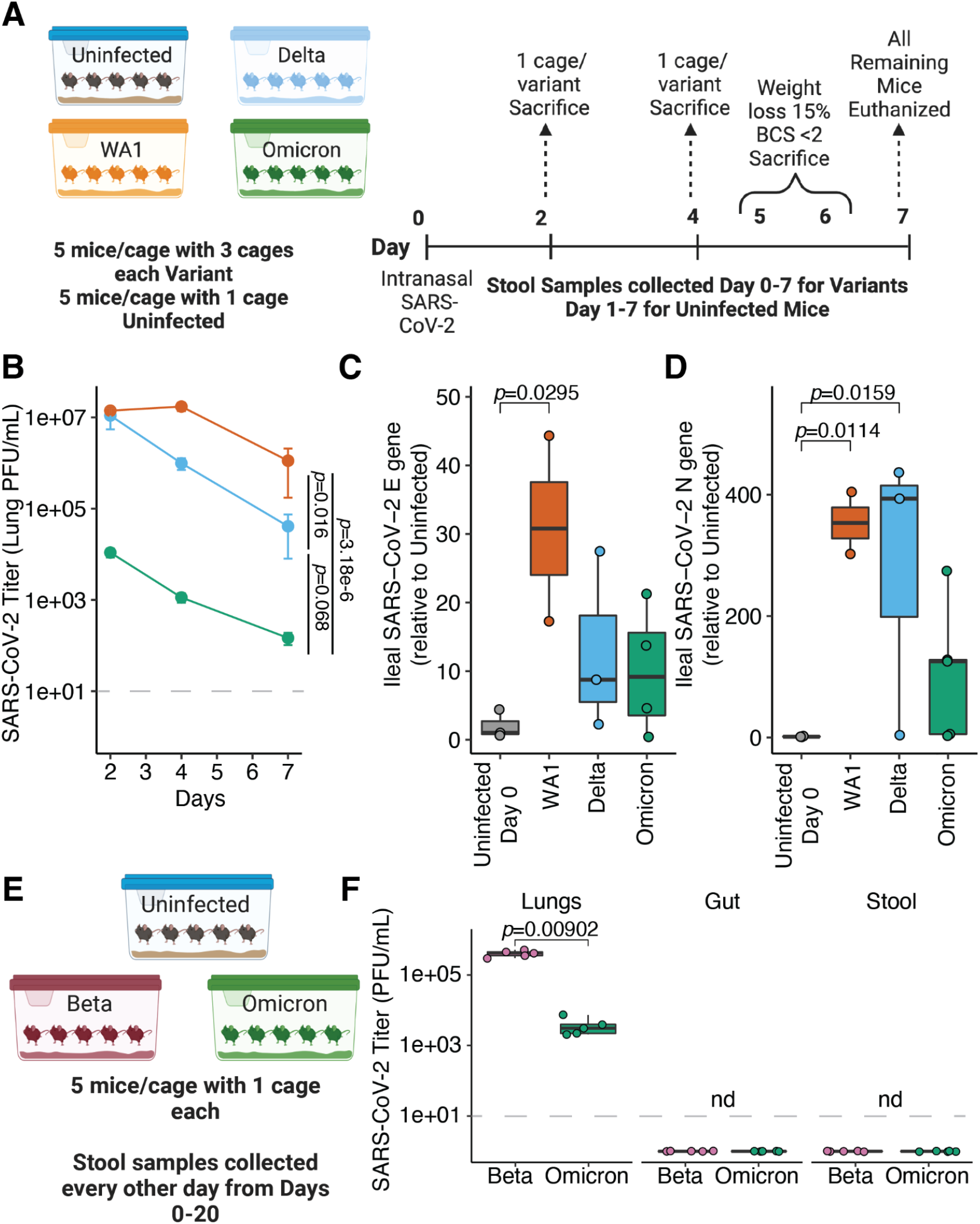
The ABSL3 SARS-CoV-2 infection model and associated viral titers. **(A)** 50 female mice were housed at a ratio of 5 mice/cage in a ABSL3 facility. Three cages of mice were infected with each variant (n=45 mice, n=15 per variant) and one cage of mice was uninfected (n=5 mice). One cage of mice was sacrificed for each variant at days 2, 4, and 7 for lung viral titer assessment. Any mice reaching a sacrificial endpoint (weight loss of 15% or body conditioning scale (BCS) < 2) were euthanized and removed from the experiment on days 5 and 6. All remaining mice were sacrificed on day 7. **(B)** Lung viral titers for each variant are shown as described in (23) with mean +/-sem plotted at each time point. Statistics represent 2 way-ANOVA and where titer is a function of variant and time, and the *p*-value for variant between the indicated comparisons is displayed. The limit of detection (LOD) for viral plaques is indicated by a gray, dashed line (LOD=10 plaque forming units (PFU) per mL). SARS-CoV-2 envelope (E) gene qPCR **(C)** or nucleocapsid (N) gene qPCR **(D)** was conducted on tissue from animals harvested at day 7 post infection. The unadjusted Dunn’s post-test of Kruskal Wallis analysis of variance is displayed for the groups with *p*<0.05 and all non-displayed comparisons are not significant. Uninfected mice were sacrificed and included as controls; given they were never infected with SARS-CoV-2 a day of infection of 0 was assigned in the figure. For **(A)** n=10 unique cages with 55 total samples from 50 mice. In **(B)** n=12 mice for WA1, n=13 mice for Delta, and n=15 mice for Omicron infected mice. In **(C-D)** each data point reflects one mouse (n=2-4 mice per group and time point). **(E)** Experimental design for C57BL/6J background infected mice. **(F)** PFUs from indicated body sites (n=5 mice per group) and statistics reflect Mann-Whitney U test between groups for lung samples. Live virus was not detected (nd) from Gut and Stool. LOD=10 PFU per mL and is indicated by a gray, dashed line.

**FIG. S7.**
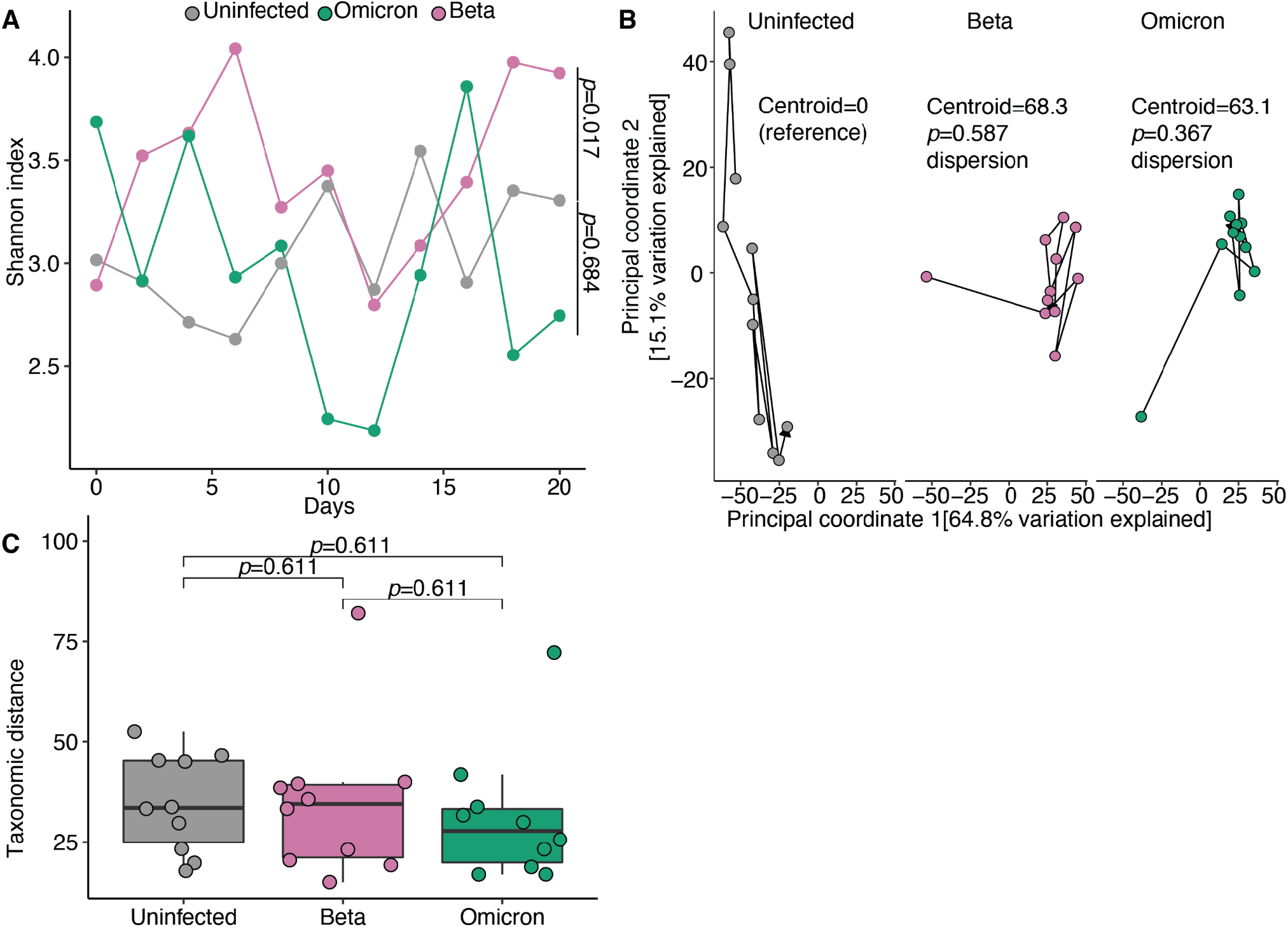
Minimal changes to alpha diversity or markers of dispersion in C57BL/6J infected mice. **(A)** Alpha diversity is plotted over time with *p*-values annotated showing the statistics for a two way ANOVA comparing the indicated variant to the uninfected group. **(B)** Principal coordinates 1 and 2 of 16S-Seq data for all analyzed samples derived from the same distance matrix and separated by group to facilitate visualization. The distance from the Uninfected reference group’s centroid is displayed for the right three panels. The *p*-value for β-dispersion from the vegan package for COVID Status is displayed and compares the variant to the uninfected reference group and adjusted using Tukey’s multiple comparisons. **(C)** Subsequent distances between successive points in **(B)** for a given cage are plotted and grouped by variants of concern. Dunn’s post-hoc test of Kruskal-Wallis analysis of variance is displayed and adjusted with a Benjamini Hochberg correction. n=15 mice with 5 mice per group in 3 cages and 33 total longitudinally collected samples.

